# xenoGI: reconstructing the history of genomic island insertions in clades of closely related bacteria

**DOI:** 10.1101/188599

**Authors:** Eliot C Bush, Anne E Clark, Carissa A DeRanek, Alexander Eng, Juliet Forman, Kevin Heath, Alexander B Lee, Daniel M Stoebel, Zunyan Wang, Matthew Wilber, Helen Wu

## Abstract

**Background:** Genomic islands play an important role in microbial genome evolution, providing a mechanism for strains to adapt to new ecological conditions. A variety of computational methods, both genome-composition based and comparative have been developed to identify them. Some of these methods are explicitly designed to work in single strains, while others make use of multiple strains. In general, existing methods do not identify islands in the context of the phylogeny in which they evolved. Even multiple strain approaches are best suited to identifying genomic islands that are present in one strain but absent in others. They do not automatically recognize islands which are shared between some strains in the clade or determine the branch on which these islands inserted within the phylogenetic tree.

**Results:** We have developed a software package, xenoGI, that identifies genomic islands and maps their origin within a clade of closely related bacteria, determining which branch they inserted on. It takes as input a set of sequenced genomes and a tree specifying their phylogenetic relationships. Making heavy use of synteny information, the package builds gene families in a species-tree-aware way, and then attempts to combine into islands those families whose members are adjacent and whose most recent common ancestor is shared. The package provides a variety of text-based analysis functions, as well as the ability to export genomic islands into formats suitable for viewing in a genome browser. We demonstrate the capabilities of the package with several examples from enteric bacteria, including an examination of the evolution of the acid fitness island in the genus Escherichia. In addition we use output from simulations and a set of known genomic islands from the literature to show that xenoGI can accurately identify genomic islands and place them on a phylogenetic tree.

**Conclusions:** xenoGI is an effective tool for studying the history of genomic island insertions in a clade of microbes. It identifies genomic islands, and determines which branch they inserted on within the phylogenetic tree for the clade. Such information is valuable because it helps us understand the adaptive path that has produced living species. Given the large and growing number of sequenced microbial genomes, this sort of analysis will become increasingly useful in the future.

## Background

Genomic islands (GI) are clusters of genes that have entered a genome via horizontal gene transfer, that is, outside the normal process of parent-offspring inheritance. Early observations were made in the context of bacterial pathogenicity, where it was found that the difference between pathogenic and non-pathogenic strains often depended on the presence of one or more genomic islands [1]. It soon became clear however, that the function of genomic islands is not restricted to pathogenicity, and that they play a broad role in microbial genome evolution [2, 3, 4].

Because of their importance, a significant number of computational methods have been developed for finding GIs. These are distinct from, but fit into a larger literature on finding individual horizontally transferred genes. GI-finding methods can be broadly divided into those that operate on a genome from a single species, and comparative genomics methods that operate on genomes from several species [5, 6].

Many single genome methods are compositional, making use of various attributes of sequence composition such as GC content, oligonucleotide frequency or codon bias. Because genomes differ in these compositional characteristics, when a foreign piece of DNA arrives into a genome, it may differ in some of these characteristics from the genome it is entering. For insertion events that are sufficiently recent, this can be a mechanism to identify foreign DNA. Such methods have been developed to try to take advantage of many compositional features, including GC content [7], oligonucleotide frequencies [8, 9, 10, 11, 12, 13, 14], and codon bias [15, 16]. Single genome methods also sometimes target specific sequence features that are associated with GI insertion such as tRNA genes [17]. And a number of such methods use combinations of multiple attributes including composition and/or specific sequence features [18, 19, 20, 21, 22, 23, 24, 25, 26].

The basic idea of comparative genomics methods is to compare related genomes to identify regions that are unique to certain genomes and likely result from horizontal transfer. These methods are closely related to methods for identifying the core and pan genomes of a set of species or reconstructing ancestral gene order [27, 28, 29, 30, 31, 32]. Comparative methods typically involve whole genome or protein alignments and then some methods built on top of this to identify orthologs and recognize events such as horizontal transfer, deletion, and so on.

Several automated comparative genomics methods for finding GIs have been developed to date. The tRNAcc package combines comparative genomics with a feature specific search [33]. It identifies islands that have inserted near tRNA genes by creating alignments between closely related species using MAUVE [34], and then looking for regions of DNA that are unique to one species near tRNA genes. This approach is good at finding those GIs that insert near tRNA genes, but will miss others. It is included in the web-based MobilomeFINDER service [21].

Another widely used method is IslandPick [35] which has been incorporated into the web service IslandViewer [36, 37, 38]. IslandPick is provided with a single input genome where the user desires to find genomic islands. It first identifies a set of comparison genomes, then creates pairwise whole genome alignments using MAUVE, and finally analyzes the alignments to identify regions that are unique in the input genome. This comparative approach allows accurate identification of GIs that are unique to the input genome, and is widely used as a part of the IslandViewer website.

Given continuing reductions in the cost of sequencing, in the future comparative genomics methods are likely to be increasingly important for finding GIs. However existing methods have one important limitation: they are not able to automatically place GIs in the context of a phylogenetic tree. Existing methods such as tRNAcc and IslandPick make use of multiple genomes, but they are best at identifying regions which are unique to one genome compared with others. If we want to study the history of genomic island insertions in a clade of microbes, these methods allow us to find islands that are unique to various strains of the clade. But they do not automatically identify genomic islands which are shared between some strains in the clade, and they do not determine the branch on which those islands inserted within the phylogenetic tree.

Here we describe xenoGI, a system that identifies genomic islands, and maps their origin within a clade of closely related bacteria. Every gene present in a clade has one of two possible origins. Either it originated in the most recent common ancestor of the clade, or it originated in a subsequent horizontal transfer event. The goal of xenoGI is to group genes by origin, identifying islands of genes that entered via common horizontal transfer events, and mapping those events onto the phylogenetic tree. Such information is often of interest because it helps us understand the adaptive path that has produced living species.

## Implementation

xenoGI is a command line program implemented in Python. It can be downloaded from https://github.com/ecbush/xenoGI.

### Input, output, basic structure

Input consists of a set of sequenced genomes in GenBank format, and a tree specifying their phylogenetic relationships. The GenBank files provide protein sequences and their genomic order in each strain. Because the algorithm makes use of synteny information, the genomes need to come from a clade of bacteria that are closely related enough to preserve gene order. For the same reason, the genome assemblies should be at the scaffold level or better. The algorithm is not suitable for analyzing plasmid sequences because of the rapid rate of change of gene content and order on plasmids. Typically the set of input genomes would include a focal clade that we wish to study, and one or two outgroups (Figure 1). These outgroups help us to better recognize core genes given the possibility of deletion in some lineages. xenoGI also includes optional scripts to help users obtain multiple alignments. These can then be used with existing methods to reconstruct a phylogenetic tree for a set of strains, if that tree is unknown.

**Figure 1.**
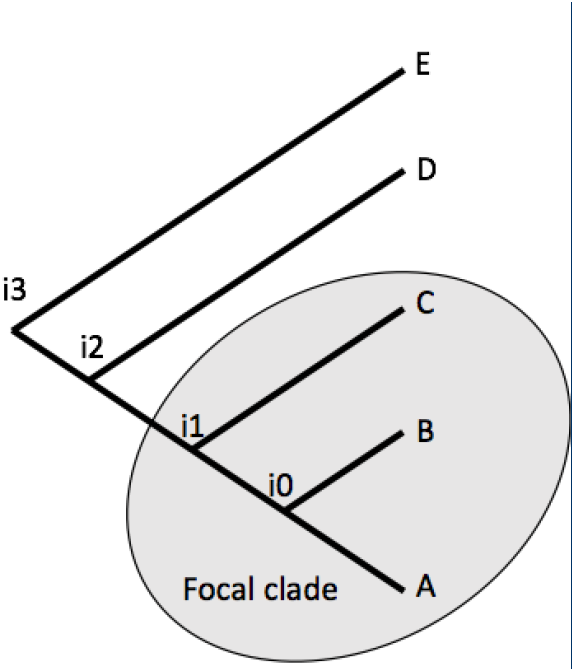
Example species tree. An input tree consisting of a focal clade and several outgroups.

Every gene in the input genomes must have one of two origins. Either it is a core gene present in the most recent common ancestor of the strains, or it arrived via a horizontal transfer event. The goal of the algorithm is to determine this origin for each gene, grouping genes that arrived together in the same horizontal transfer events as islands. The output is a text file specifying these islands. The output can be visualized further with several text-based visualization functions included in the package, and can also be exported for visualization in a genome browser.

There are three basic steps the algorithm takes. It first calculates a set of scores between genes in the input genomes based on their protein sequences. This includes scores based on sequence similarity and scores based on synteny. It next groups genes into families in a tree-aware way. Finally it groups these families into islands where the families in an island are interpreted to have a common origin (they either arrived in the same horizontal transfer event or are core genes).

Every gene family thus formed has a most recent common ancestor that falls on some node in the input tree.

### Calculating scores between genes

The first step is to calculate a set of similarity and synteny scores between genes in the input data set. We initially run protein BLAST between every gene and genes in all the other strains. For those pairs of genes above a certain E-value threshold, we calculate a number of other types of scores.

A raw score is a similarity score between a pair of proteins. We take the global alignment score and scale it to be between 0 and 1, using the following formula:

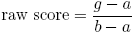

where *g* is the global alignment score between two proteins. *a* is a floor value for the alignment score between these two proteins based on what we would get if they were aligned with all gaps (among the pairs we look at, which have significant BLAST hits, there will be nothing lower than this). *b* is a ceiling value for the alignment score (the score of the shorter aligned against itself).

The global alignment is calculated using Parasail [39]. The use of global alignment here reflects the fact that we are operating in a clade of closely related strains and the gene families we build consist of closely related genes. Because of this, we expect alignments between homologs within families to span entire proteins making global alignment preferable to local.

The calculation of raw scores can be run in parallel on multiple processors.

We also calculate a normalized similarity score, which normalizes for the average level of protein distance between a pair of species. Such scores make it easier to set thresholds based on similarity in family formation.

To begin, we identify sets of orthologs where there is exactly one copy in each strain. We do this with the all around best reciprocal hit method, identifying sets of orthologs where every gene is a best reciprocal hit with every other gene, and has one copy in each strain. These sets of orthologs are very conservative and high confidence. We’ll refer to them below as conservative core genes.

Then for each pair of strains we calculate the mean and standard deviation of raw scores between all pairs of orthologs in these sets of conservative core genes. Using this, we take a raw score comparing proteins in two strains and normalize it as follows:

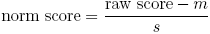

where *m* is the mean and *s* is the standard deviation of raw scores from the conservative core genes for that pair of strains.

We calculate two kinds of synteny scores based on similarity in the neighborhood of genes.

Our core synteny score makes use of the conservative core genes just discussed. To calculate this score, we define a neighborhood for each gene consisting of a user specified window of conservative core genes in either direction. For example, the neighborhood might be a region encompassing 20 conservative core genes, the first 10 in either direction. To compare a gene A from one strain with a gene B from another, we determine how many conservative core genes from the neighborhood of A are also found in the neighborhood of B, and divide this by the number of conservative core genes in the neighborhood (twenty in the example above). This core-gene synteny score is then:

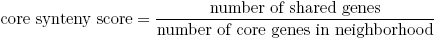

and is thus a value between 0 and 1.

Because the conservative core gene neighborhoods tend to be large, this measure of synteny looks at a comparatively large region around a gene.

To calculate synteny in a more local region, we also calculate a synteny score based on all genes, including non-core genes.

To compare a gene A from one strain with a gene B from another, we obtain lists of neighboring genes for each. Let *N*_*a*_ be the set of neighboring genes for gene A, taken from within a window of size *W* measured in number of genes, and including both core and non-core genes. Let *N*_*b*_ be the same for gene B. Our local synteny score is calculated as follows. We find the pair of genes with the highest norm score between *N*_*a*_ and *N*_*b*_ and keep it. We remove those genes from *N*_*a*_ and *N*_*b*_, find the next highest pair between them and so on. The local synteny score for gene A and gene B is the average of the *T* highest scoring pairs.

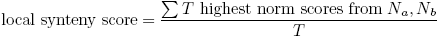

The calculation of synteny scores can be run in parallel on multiple processors.

### Forming gene families in a tree-aware way

We wish to build gene families that fit into the known species tree.

For our purposes, a gene family is a set of genes that have descended from a single ancestor gene that existed within that species tree. The most recent common ancestor (MRCA) in such a gene family will fall on the species tree, and its location there will reflect the origin of the family. Gene families whose MRCA falls outside the focal clade are core genes relative to that clade. Gene families whose MRCA falls within the focal clade have arrived via horizontal transfer. This implies that genes that share a deeper homology predating the root of the species tree, will be placed in different families. In cases where the same gene has entered our clade multiple times as a part of distinct transfer events, we want each insertion to correspond to a different gene family.

Our approach is to use a version of the PHiGs algorithm [40] which we have modified to consider synteny information. The PhIGs algorithm operates on the species tree beginning at the root node and moving successively to descendant nodes. At each node, we group all genes descending from species on the left branch (call this group 1), and also group all genes descending from species on the right branch (call this group 2). For nodes other than the root, there are also genes in outgroup species, which we ignore. For every gene in one group, we find the most similar gene in the other, e.g. for all genes in group 1 we find the closest gene in group 2. A pair identified in this way constitutes a seed upon which we build a larger family via single linkage clustering. Because order matters for this approach, the seeds are processed in order of similarity, so that the most similar seeds are worked on first. To build a family from a seed, we identify all previously unclustered genes in either group 1 or group 2 that are closer to a member of the growing family then the original seeds in 1 and 2 were to each other. In our implementation, similarity is measured by the raw score.

We have modified the original algorithm in several ways. In order to add a gene to a family, the basic algorithm requires that it be more similar to some family member than the similarity level of the seed. We have added an absolute threshold for similarity measured using the norm score. This is typically set low, and is a sanity check to make sure we’re not clustering very distantly related proteins together into a family. We also incorporate synteny in a number of ways. We have synteny thresholds for both core and local synteny. Below these thresholds, we do not add a gene to a family. We also use high synteny on the other end, to increase the raw score between a pair of genes we’re considering, potentially helping them get over the bar of seed similarity set by the basic algorithm.

This step runs on a single processor.

### Grouping families into islands

Our goal is to group gene families that arrived together as a part of the same horizontal transfer event. We refer to such groups as islands.

We first sort families according to their MRCA because families that belong in the same island will have the same MRCA. We then build islands using a greedy approach that progressively adds families that are inferred to be adjacent to each other in the MRCA.

We identify genes to add using a parsimony-based metric. Our metric uses a very simple notion of evolutionary changes in gene order. If a pair of genes were adjacent, but due to rearrangements move apart, we assess a cost of one. Similarly if two genes were non-adjacent, but due to rearrangements move to be adjacent, we also assess a cost of one. Consider two gene families with the same MRCA, for example i0 in Figure 1. We have adjacency information for those families in species A and species B. Our approach is to calculate a rearrangement cost for these families under two cases: either assuming they were adjacent at their MRCA, or assuming they were not. We define the rearrangement score for the families as follows:

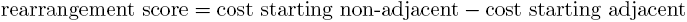

This score can range between -2 and 2 with more positive values indicating families that are more likely to have been adjacent in the common ancestor.

We begin by creating a set of one-family islands for all families with a particular MRCA. We then calculate the rearrangement score for every pair of islands and identify the pair with the highest score, arbitrarily breaking ties. Then we merge this pair into a two-family island and recalculate its rearrangement score with other islands. Note that because multi-family islands have a gene order (the inferred order in the MRCA), to calculate rearrangement scores for them, we consider adjacency on each of their two ends. The algorithm continues merging islands until all the rearrangement scores are below a certain threshold. It then repeats the procedure relaxing the criterion for adjacency (e.g. we can count as adjacent genes that are one gene away from each other). When all the rearrangement scores are below threshold, the algorithm terminates.

The island-making step runs on multiple processors.

### Analysis tools

Included in the package are command line tools for identifying and visualizing islands at particular nodes, for finding islands associated with particular genes and for examining gene families. Also included are scripts for exporting xenoGI island output to bed or gff format for visualization in a genome browser such as IGB [41].

## Results

### Validation via simulation

We assessed the effectiveness of xenoGI in two ways: using simulations, and detecting known genomic islands.

We used simulations to produce test data sets where the location of GI insertions within a phylogenetic tree was known. To do this we implemented a custom simulator that evolved sequences over a user provided phylogenetic tree, allowing for horizontal transfer of novel genes (from outside the clade) as well as for genomic scale deletions, duplications and inversions, and amino acid level sequence change. The latter was done using the pyvolve module [42].

Figure 2 shows the results of a simulation on a tree with 11 species. The simulation was run on a tree matching the branch lengths and topology of a tree from the enteric bacteria, discussed below. Species A-I form the focal clade, with J and K as outgroups. The simulation contained a total of 495 horizontal transfer events which mapped onto the various branches. These ranged in size from 2 to 51 genes. There were also 1009 deletions (from 1 to 50 genes in size), 494 duplications (from 1 to 48 genes in size), and 125 inversions (from 5 to 147 genes in size).

**Figure 2.**
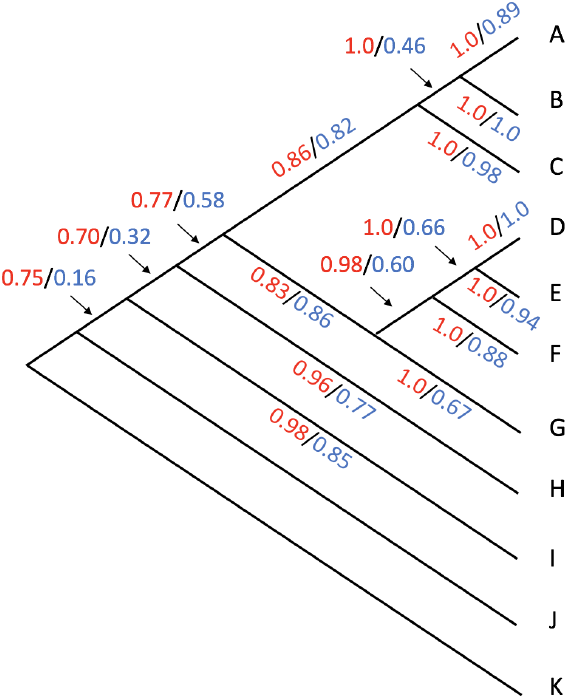
Phylogenetic tree used in genome simulations. We ran xenoGI on simulated genomes that were generated on the tree shown. On each branch we show the true positive rate (red) and the positive predictive value (blue) for xenoGI on that branch.

xenoGI achieved a base-wise true positive rate of 0.85 and a positive predictive value of 0.51 over the whole tree. What we mean by base-wise, taking the example of the true positive rate, is that among all nucleotides that were truly in genomic islands, xenoGI correctly identified (and placed on the correct branch) 0.85 of them. The majority of islands (0.83) had just a single xenoGI predicted island overlapping them. As can be seen from the figure, xenoGI’s accuracy is best for GIs inserting on tips, and declines as we move to internal branches.

### Validation via GIs from the literature

Simulations have the advantage of providing a situation where ground truth is known. However they are necessarily simplified, and may not adequately capture the features of real genome evolution. For this reason, we also examined how well xenoGI identified genomic islands that have been reported in the literature.

Wei et al. compiled a list of known genomic islands from 13 bacterial strains for the purposes of validation [26]. We used six of these for training and development, and saved five for the validation which we report here. For each of the five strains, we identified four or more closely related species with genomes in genbank. We reconstructed the history of genomic island insertions in the resulting clade using the default parameters we had developed working with the first six. A listing of the strains and trees used here can be found in Additional File 1. We note that there were not many cases where these known islands were shared among multiple already sequenced genomes. Thus most of the cases we look at here, the islands are on the tips of the phylogenies we created. In the section below we give several examples of cases where islands can be identified on internal branches of a tree.

The islands in this validation set were reported in genome papers for their respective strains [43, 44, 45, 46, 47]. They are chiefly based on comparative work involving human curation, and in some cases on experimental evidence as well. They are likely to represent true islands. However we cannot be sure that these represent exhaustive lists of all genomic islands in a strain. For this reason, it does not make sense to calculate a true positive rate or positive predictive value here. The validation ranges are given in nucleotide positions, whereas xenoGI identifies genes that are part of an island. For the purposes of comparison, we consider the nucleotides of a xenoGI island to be those between the beginning of the first gene and the end of the last gene.

Table 1 shows our results for this analysis. As can be seen from the table, the base coverage, that is the proportion of validation island bases that are covered by a xenoGI island, is in the upper 90 percent range for 4/5 strains, and 88% for Cronobacter. In addition, in the majority of cases, xenoGI identified a single island corresponding to each validation island. And in most cases (6/11) where a validation island was broken into more than one xenoGI island, this was actually correct, due to the fact that the validation island had genes with a most recent common ancestor at different times on the tree, and likely resulted from multiple horizontal transfer events.

**Table 1.** Summary of xenoGI results on validation cases in several strains. Each row corresponds to a strain. Num. islands represents the number of validation islands and total bases represents the total number of nucleotides in those islands. Base coverage is the proportion of all bases in the validation islands that xenoGI correctly recognized as an island. In single island indicates the proportion of validation islands that xenoGI captured as a single island.

|  | Num. islands | Total bases | Base coverage | In single island |
| --- | --- | --- | --- | --- |
| *Burkholderia cenocepacia* J2315 | 13 | 530,772 | 0.977 | 0.917 |
| *Corynebacterium diphtheriae* NCTC 13129 | 13 | 249,918 | 0.983 | 0.769 |
| *Cronobacter sakazakii* ATCC BAA-894 | 14 | 305,124 | 0.879 | 0.643 |
| *Streptococcus equi* 4047 | 7 | 243,337 | 0.951 | 0.857 |
| *Vibrio cholerae* O1 biovar eltor str. N16961 | 6 | 295,096 | 0.970 | 0.833 |

Additional file 1 contains a more detailed description of our comparison between xenoGI islands and validation islands. One of the measures included is extra coverage, which refers to cases where a xenoGI island extends beyond the range of the validation island. There were 21 cases where the extra coverage was more than 10% of the length of the island. In the majority of these (16/21) further examination suggested that in fact xenoGI was correct to extend the range of the island.

### Reconstructing the timing of GI insertions: several examples from enteric bacteria

We look at two examples of genomic islands from the enteric bacteria with the goal of demonstrating the sort of analysis one can do with xenoGI. We do this on a tree of eleven enteric species (Figure 3 A).

**Figure 3.**
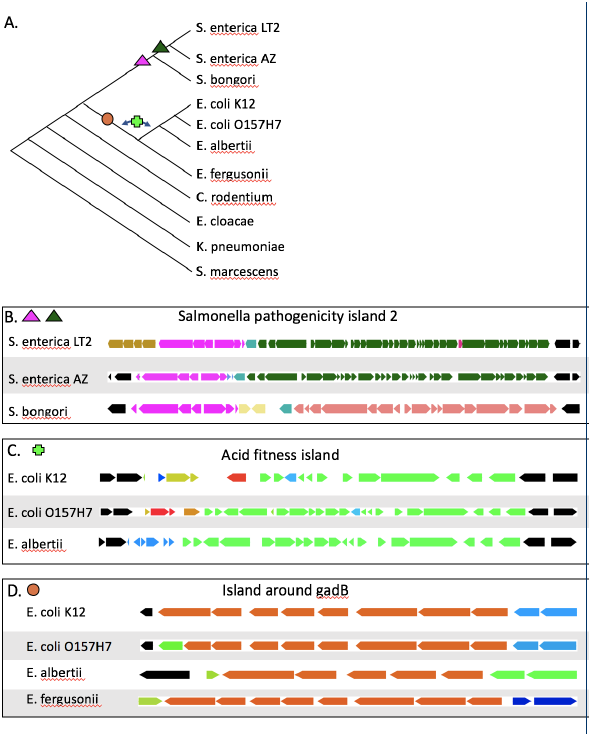
Examples from enteric bacteria. A. Phylogenetic tree of 11 enteric species. Symbols indicate the branches of insertion of GIs in B-D. The images in B-D were made by outputting xenoGI islands and then displaying in the IGB genome browser. Note that the scale for the three is not exactly the same. In the figures, different islands are given different colors. All islands with an mrca at or before the point where *C. rodentium* diverges are colored black. B. Salmonella pathogenicity island 2 shown in three *Salmonella* species. C. The acid fitness island as reconstructed by xenoGI in two *E. coli* species and *E. albertii*. D. The island around gadB in our four *Escherichia* species.

The first example is Salmonella pathogenicity island 2 (SPI-2), from *Salmonella enterica* [48, 49]. This island is essential for virulence in *S. enterica* and contains a type-III secretion system that is expressed while the bacterial cell is inside host cells [50]. The island was originally defined as lying between the genes ydhE and pykF [51]. It is known to consist of several components with different evolutionary origins [52].

xenoGI’s results are consistent with what we would expect from the literature. The largest part of SPI-2 is colored dark green in Figure 3 B. It includes the type III secretion system and is shared between *Salmonella enterica* strains (LT2 and Arizoniae in our sample), but is not present in *Salmonella bongori*. xenoGI puts this region in one island, reflecting the common origin of the genes. All the genes in the region are included, with the exception of one gene specific to *S. enterica LT2* which is shown in red. xenoGI identifies another island (shown in pink) corresponding to a second part of SPI-2. This region contains the tetrathionate reductase gene cluster. Consistent with the literature, xenoGI identifies this region as being shared between *S. enterica* and *S. bongori* [52]. The final part of SPI-2, shown in brown in Figure 3B, contains genes that are present in *S. enterica LT2*, but not *S. enterica arizoniae*.

Our second example is the acid fitness island (AFI) in *E. coli* [53]. This island runs from slp to gadA (Figure 3C) in *E. coli K12* and is about 15 kb long [54]. It encodes a number of genes involved in resistance to acid stress including a glutamate decarboxylase enzyme (coded for by gadA) and its regulators. xenoGI identified the island between slp and gadA, and placed it on the branch after the divergence of *E. fergusonii*, but before the divergence of *E. albertii*. One internal gene which is part of the island, yhiD, was left out because it is not present in *E. albertii*.

Further exploration reveals some additional features of the evolution of acid fitness genes in *Escherichia*. The acid fitness island was originally described in *E. coli K12* [53], and contains 12 genes in that strain. However the island in both *E. coli O157H7* and *E. albertii* contains an additional 8 genes. As it turns out, these additional genes, which are involved in the scavenging of iron from hosts, have been identified and studied previously. They were identified first in *Shigella dysenteriae*, but were found to also be present in a number of *E. coli* strains [55]. (It’s worth noting that *Shigella* strains nest within the *E. coli* clade.) This so-called heme transport locus falls in the middle of the AFI, but because the AFI has mostly been studied in *E. coli K12*, and that strain lacks the heme transport locus, it was never recognized that the two are co-localized. xenoGI places them in the same island because they have the same most recent common ancestor, being present in some *E. coli* strains and *E. albertii*. It is possible that these two sets of genes with seemingly distinct functions arrived as a part of a single event.

A second question relates to the time of arrival of the AFI. xenoGI places it on the branch before *E. albertii* diverged (Figure 3A). However, an analysis of AFI homologs in other strains using several functions included in the package reveals that *E. fergusonii* contains a nearly complete copy of the AFI (missing only the heme transport genes and gadA), but in a non-syntenic location. The question then is whether this island in *E. fergusonii* represents an independent insertion. Alternatively, it may have resulted from the same insertion as in the other *Escherichia* strains, but have been moved to a different location via some rearrangement or translocational process. It has been observed that *E. fergusonii* has undergone a large number of genome rearrangements since the time of its divergence from *E. coli* [27]. That notwithstanding, the complete lack of synteny here favors an independent insertion, and is the reason xenoGI placed *E. fergusonii’s* AFI in a separate island.

Another issue is the phylogenetic distribution of the heme transport genes. The original paper on these genes observed a puzzling phylogenetic distribution in *E. coli* [55]. We observe something similar in our enteric species. The heme transport locus is not present in *E. coli K12* and *E. fergusonii*, but is present in *E. coli O157H7* and *E. albertii*. This distribution would require an insertion into and then a clean deletion from the AFI, or else some process involving horizontal transfer and perhaps gene conversion.

The status of another set of genes can also shed light on the evolution of acid tolerance in *Escherichia*. *Escherichia* genomes contain a second glutamate decarboxylase enzyme, gadB. gadB has typically been seen as the result of a gene duplication, the idea being that the AFI was inserted, and then gadB arose by duplication [56]. However xenoGI finds that the gadB gene lies in an island consisting of 8 genes that is shared by the entire *Escherichia* clade (Figure 3D), including *E. fergusonii*, but is not found outside that group. That is, it appears to have arisen on the branch leading to *Escherichia* before the divergence of *E. fergusonii*. This fact raises the question as to whether gadB is really a duplication or the result of an independent insertion event (albeit one that may have been followed by gene conversion events between gadA and gadB [56]). The other genes in the island surrounding gadB do not have close paralogs in other parts of the genome, a fact which may favor the idea of an independent insertion via horizontal transfer.

Acid tolerance has often been seen as a feature unique to *E. coli* [56], however our results show that this is in fact a characteristic of the whole *Escherichia* clade. This fact does have some practical significance, as AFI genes have been the basis for assays attempting to identify *E. coli* in samples [57, 58]. Beyond this our data suggest that the evolution of acid fitness in this group was more complicated than previously appreciated, likely involving multiple insertion events.

### Resource usage

We have run xenoGI on up to 115 strains. There is a trade off between time and RAM usage, because using more processors requires more RAM. Additional file 2 shows plots of RAM usage, user time and wall clock time for up to 40 strains running on 50 processors. On our machine, 40 strains required approximately 500GB of ram and around 20 hours from start to finish.

## Discussion

The results presented above show that xenoGI can effectively reconstruct genomic islands. In the validation using simulations, xenoGI’s true positive rate and positive predictive value were high, and most insertions were recognized as a single island. We did observe that xenoGI’s effectiveness declined as we moved to more internal branches. This trend is not surprising. Internal branches are older, and in the extra time that has passed, events may have occurred that obscure the evidence for a horizontal transfer event. xenoGI also correctly identified most of the previously identified validation islands in real genomes (Table 1). Often in cases where it seemingly made a mistake, e.g. split a single validation range into several islands, upon closer examination we found that its result was correct (Additional File 1).

It is worth reflecting briefly on some of the assumptions and limitations of our approach. xenoGI makes extensive use of synteny information. This means that it will be ineffective in situations where gene order or composition is changing rapidly, for example on plasmids which typically lack conservative core genes as a result of rapid evolution. In such cases, xenoGI will find large islands of seemingly unique genes. However these are not really unique, but only seem so because they lack enough synteny with their true homologs to be put together in families.

Along similar lines, in genomes where there has been an unusual amount of genome rearrangement, the resulting reduction in synteny makes it more challenging to use xenoGI. It is possible to adjust the parameters to compensate for this to some extent, e.g. by shrinking the synteny window sizes.

A third caveat has to do with genomic island insertion hotspots. It has been observed that certain regions are more likely than others to receive insertions [27]. When there are multiple insertions of the same or very similar GIs, xenoGI distinguishes these insertions using synteny. However if similar islands insert multiple times in the same region, it will not be able to recognize those events as distinct.

Future work might attempt to use additional information such as that available in gene trees to supplement decisions we are currently making based on synteny. This could potentially enable us to better recognize cases where related islands inserted in the same location multiple times.

More generally, the problem of making tree-aware families is one that might be aided with machine learning approaches. The creation of such families involves integrating multiple pieces of information. However it is challenging to create a single set of rules that captures what we want to do. It may be easier for a human to annotate a clade of bacterial genomes, creating a set of gene families, and then let a machine learning algorithm learn from that. The algorithm would be using similar information to our current algorithm, but would have learned the rules based on a training set.

Finally, there are a number of examples of genomic islands which have inserted multiple times in different strains. The AFI discussed above is a potential example. It would be helpful to have a more systematic way to identify this. We could potentially add an additional step which involves comparing all the islands found in a clade against each other.

## Conclusions

As more and more microbial genomes are sequenced, it becomes desirable to analyze genomic adaptation in the context of phylogenetic trees. Here we have presented xenoGI, a software package that takes a clade of closely related microbes and identifies islands of genes that entered via common horizontal transfer events, placing those events on the phylogenetic tree for the clade.

## List of abbreviations

GI: genomic island MRCA: most recent common ancestor AFI: acid fitness island

## Funding

This work was supported by HHMI Undergraduate Science Education award number 52007544 to Harvey Mudd College.

## Availability of data and materials

xenoGI and related materials can be downloaded from https://github.com/ecbush/xenoGI.

## Competing interests

The authors declare that they have no competing interests.

## Author’s contributions

EB and DS identified the problem, and EB designed the approach. EB, AC, CD, AE, JF, KH, AL, MW and HW implemented the software. CD, DS and EB worked on testing and validation, and EB wrote the manuscript.

## Acknowledgements

We thank Yi-Chieh Wu for helpful discussions, Bingxin Lu for suggestions on validation and Stephanie Spielman for help in using pyvolve for our purposes.

## Tables

### Additional Files

Additional file 1 — A more detailed description of the correspondence between xenoGI islands and validation islands. A tab-delimited text file containing each validation range used in the five species we looked at. Includes the assemblies used to compare with each strain and the tree. For each validation range gives the base-wise coverage, the amount of extra coverage, the number of overlapping xenoGI islands and any comments.

Additional file 2 — Resource usage of xenoGI

Plots of RAM usage, user time and wall clock time for up to 40 strains running on 50 processors.

## References

1 Hacker, J., Bender, L., Ott, M., Wingender, J., Lund, B., Marre, R., Goebel, W.: Deletions of chromosomal regions coding for fimbriae and hemolysins occur in vitro and in vivo in various extra intestinal escherichia coli isolates. Microbial pathogenesis 8(3), 213–225 (1990)

2 Hacker, J., Kaper, J.B.: Pathogenicity islands and the evolution of microbes. Annual Reviews in Microbiology 54(1), 641–679 (2000)

3 Dobrindt, U., Hochhut, B., Hentschel, U., Hacker, J.: Genomic islands in pathogenic and environmental microorganisms. Nature Reviews Microbiology 2(5), 414–424 (2004)

4 Ochman, H., Lawrence, J.G., Groisman, E.A.: Lateral gene transfer and the nature of bacterial innovation. Nature 405(6784), 299–304 (2000)

5 Langille, M.G., Hsiao, W.W., Brinkman, F.S.: Detecting genomic islands using bioinformatics approaches. Nature Reviews Microbiology 8(5), 373–382 (2010)

6 Lu, B., Leong, H.W.: Computational methods for predicting genomic islands in microbial genomes. Computational and structural biotechnology journal 14, 200–206 (2016)

7 Zhang, R., Zhang, C.-T.: A systematic method to identify genomic islands and its applications in analyzing the genomes of corynebacterium glutamicum and vibrio vulnificus cmcp6 chromosome i. Bioinformatics 20(5), 612–622 (2004)

8 Sandberg, R., Winberg, G., Br¨anden, C.-I., Kaske, A., Ernberg, I., Cöster, J.: Capturing whole-genome characteristics in short sequences using a naive bayesian classifier. Genome research 11(8), 1404–1409 (2001)

9 Tsirigos, A., Rigoutsos, I.: A new computational method for the detection of horizontal gene transfer events. Nucleic acids research 33(3), 922–933 (2005)

10 Vernikos, G.S., Parkhill, J.: Interpolated variable order motifs for identification of horizontally acquired dna: revisiting the salmonella pathogenicity islands. Bioinformatics 22(18), 2196–2203 (2006)

11 Rajan, I., Aravamuthan, S., Mande, S.S.: Identification of compositionally distinct regions in genomes using the centroid method. Bioinformatics 23(20), 2672–2677 (2007)

12 Chatterjee, R., Chaudhuri, K., Chaudhuri, P.: On detection and assessment of statistical significance of genomic islands. BMC genomics 9(1), 150 (2008)

13 Arvey, A.J., Azad, R.K., Raval, A., Lawrence, J.G.: Detection of genomic islands via segmental genome heterogeneity. Nucleic acids research 37(16), 5255–5266 (2009)

14 Shrivastava, S., Reddy, C.V.S.K., Mande, S.S.: Indegenius, a new method for high-throughput identification of specialized functional islands in completely sequenced organisms. Journal of biosciences 35(3), 351–364 (2010)

15 Merkl, R.: Sigi: score-based identification of genomic islands. BMC bioinformatics 5(1), 22 (2004)

16 Waack, S., Keller, O., Asper, R., Brodag, T., Damm, C., Fricke, W.F., Surovcik, K., Meinicke, P., Merkl, R.: Score-based prediction of genomic islands in prokaryotic genomes using hidden markov models. BMC bioinformatics 7(1), 142 (2006)

17 Hudson, C.M., Lau, B.Y., Williams, K.P.: Islander: a database of precisely mapped genomic islands in trna and tmrna genes. Nucleic acids research 43(D1), 48–53 (2015)

18 Karlin, S.: Detecting anomalous gene clusters and pathogenicity islands in diverse bacterial genomes. Trends in microbiology 9(7), 335–343 (2001)

19 Tu, Q., Ding, D.: Detecting pathogenicity islands and anomalous gene clusters by iterative discriminant analysis. FEMS microbiology letters 221(2), 269–275 (2003)

20 Hsiao, W.W., Ung, K., Aeschliman, D., Bryan, J., Finlay, B.B., Brinkman, F.S.: Evidence of a large novel gene pool associated with prokaryotic genomic islands. PLoS Genet 1(5), 62 (2005)

21 Ou, H.-Y., He, X., Harrison, E.M., Kulasekara, B.R., Thani, A.B., Kadioglu, A., Lory, S., Hinton, J.C., Barer, M.R., Deng, Z., et al.: Mobilomefinder: web-based tools for in silico and experimental discovery of bacterial genomic islands. Nucleic acids research 35(suppl 2), 97–104 (2007)

22 Pundhir, S., Vijayvargiya, H., Kumar, A.: Predictbias: a server for the identification of genomic and pathogenicity islands in prokaryotes. In silico biology 8(3, 4), 223–234 (2008)

23 Wei, W., Guo, F.: Prediction of genomic islands in seven human pathogens using the z-island method. Genet. Mol. Res 10, 2307–2315 (2011)

24 Soares, S., Abreu, V., Ramos, R., Cerdeira, L., Silva, A., et al.: Pips: Pathogenicity island prediction software. PLoS ONE 7(2), 30848 (2012)

25 Lee, C.-C., Chen, Y.-P.P., Yao, T.-J., Ma, C.-Y., Lo, W.-C., Lyu, P.-C., Tang, C.Y.: Gi-pop: a combinational annotation and genomic island prediction pipeline for ongoing microbial genome projects. Gene 518(1), 114–123 (2013)

26 Wei, W., Gao, F., Du, M.-Z., Hua, H.-L., Wang, J., Guo, F.-B.: Zisland explorer: detect genomic islands by combining homogeneity and heterogeneity properties. Briefings in bioinformatics 18(3), 357–366 (2017)

27 Touchon, M., Hoede, C., Tenaillon, O., Barbe, V., Baeriswyl, S., Bidet, P., Bingen, E., Bonacorsi, S., Bouchier, C., Bouvet, O., et al.: Organised genome dynamics in the escherichia coli species results in highly diverse adaptive paths. PLoS genet 5(1), 1000344 (2009)

28 Laing, C., Buchanan, C., Taboada, E.N., Zhang, Y., Kropinski, A., Villegas, A., Thomas, J.E., Gannon, V.P.: Pan-genome sequence analysis using panseq: an online tool for the rapid analysis of core and accessory genomic regions. BMC bioinformatics 11(1), 461 (2010)

29 Fouts, D.E., Brinkac, L., Beck, E., Inman, J., Sutton, G.: Panoct: automated clustering of orthologs using conserved gene neighborhood for pan-genomic analysis of bacterial strains and closely related species. Nucleic acids research 40(22), 172–172 (2012)

30 Yang, K., Heath, L.S., Setubal, J.C.: Regen: Ancestral genome reconstruction for bacteria. Genes 3(3), 423–443 (2012)

31 Contreras-Moreira, B., Vinuesa, P.: Get homologues, a versatile software package for scalable and robust microbial pangenome analysis. Applied and environmental microbiology 79(24), 7696–7701 (2013)

32 Paul, S., Bhardwaj, A., Bag, S.K., Sokurenko, E.V., Chattopadhyay, S.: Pancoregen—profiling, detecting, annotating protein-coding genes in microbial genomes. Genomics 106(6), 367–372 (2015)

33 Ou, H.-Y., Chen, L.-L., Lonnen, J., Chaudhuri, R.R., Thani, A.B., Smith, R., Garton, N.J., Hinton, J., Pallen, M., Barer, M.R., et al.: A novel strategy for the identification of genomic islands by comparative analysis of the contents and contexts of trna sites in closely related bacteria. Nucleic acids research 34(1), 3–3 (2006)

34 Darling, A.C., Mau, B., Blattner, F.R., Perna, N.T.: Mauve: multiple alignment of conserved genomic sequence with rearrangements. Genome research 14(7), 1394–1403 (2004)

35 Langille, M.G., Hsiao, W.W., Brinkman, F.S.: Evaluation of genomic island predictors using a comparative genomics approach. BMC bioinformatics 9(1), 329 (2008)

36 Langille, M.G., Brinkman, F.S.: Islandviewer: an integrated interface for computational identification and visualization of genomic islands. Bioinformatics 25(5), 664–665 (2009)

37 Dhillon, B.K., Chiu, T.A., Laird, M.R., Langille, M.G., Brinkman, F.S.: Islandviewer update: improved genomic island discovery and visualization. Nucleic acids research, 394 (2013)

38 Bertelli, C., Laird, M.R., Williams, K.P., Lau, B.Y., Hoad, G., Winsor, G.L., Brinkman, F.S.: Islandviewer 4: expanded prediction of genomic islands for larger-scale datasets. Nucleic Acids Research

39 Daily, J.: Parasail: Simd c library for global, semi-global, and local pairwise sequence alignments. BMC bioinformatics 17(1), 81 (2016)

40 Dehal, P.S., Boore, J.L.: A phylogenomic gene cluster resource: the phylogenetically inferred groups (phigs) database. BMC bioinformatics 7(1), 201 (2006)

41 Freese, N.H., Norris, D.C., Loraine, A.E.: Integrated genome browser: visual analytics platform for genomics. Bioinformatics 32(14), 2089–2095 (2016)

42 Spielman, S.J., Wilke, C.O.: Pyvolve: a flexible python module for simulating sequences along phylogenies. PloS one 10(9), 0139047 (2015)

43 Heidelberg, J.F., Eisen, J.A., Nelson, W.C., Clayton, R.A., Gwinn, M.L., Dodson, R.J., Haft, D.H., Hickey, E.K., Peterson, J.D., Umayam, L., et al.: Dna sequence of both chromosomes of the cholera pathogen vibrio cholerae. Nature 406(6795), 477–483 (2000)

44 Cerde∼no-T´arraga, A., Efstratiou, A., Dover, L., Holden, M., Pallen, M., Bentley, S., Besra, G., Churcher, C., James, K., De Zoysa, A., et al.: The complete genome sequence and analysis of corynebacterium diphtheriae nctc13129. Nucleic acids research 31(22), 6516–6523 (2003)

45 Holden, M.T., Seth-Smith, H.M., Crossman, L.C., Sebaihia, M., Bentley, S.D., Cerdeño-Tárraga, A.M., Thomson, N.R., Bason, N., Quail, M.A., Sharp, S., et al.: The genome of burkholderia cenocepacia j2315, an epidemic pathogen of cystic fibrosis patients. Journal of bacteriology 191(1), 261–277 (2009)

46 Holden, M.T., Heather, Z., Paillot, R., Steward, K.F., Webb, K., Ainslie, F., Jourdan, T., Bason, N.C., Holroyd, N.E., Mungall, K., et al.: Genomic evidence for the evolution of streptococcus equi: host restriction, increased virulence, and genetic exchange with human pathogens. PLoS pathogens 5(3), 1000346 (2009)

47 Kucerova, E., Clifton, S.W., Xia, X.-Q., Long, F., Porwollik, S., Fulton, L., Fronick, C., Minx, P., Kyung, K., Warren, W., et al.: Genome sequence of cronobacter sakazakii baa-894 and comparative genomic hybridization analysis with other cronobacter species. PloS one 5(3), 9556 (2010)

48 Ochman, H., Soncini, F.C., Solomon, F., Groisman, E.A.: Identification of a pathogenicity island required for salmonella survival in host cells. Proceedings of the National Academy of Sciences 93(15), 7800–7804 (1996)

49 Shea, J.E., Hensel, M., Gleeson, C., Holden, D.W.: Identification of a virulence locus encoding a second type iii secretion system in salmonella typhimurium. Proceedings of the National Academy of Sciences 93(6), 2593–2597 (1996)

50 Figueira, R., Holden, D.W.: Functions of the salmonella pathogenicity island 2 (spi-2) type iii secretion system effectors. Microbiology 158(5), 1147–1161 (2012)

51 Hensel, M., Shea, J.E., Bäumler, A.J., Gleeson, C., Blattner, F., Holden, D.W.: Analysis of the boundaries of salmonella pathogenicity island 2 and the corresponding chromosomal region of escherichia coli k-12. Journal of bacteriology 179(4), 1105–1111 (1997)

52 Vernikos, G.S., Thomson, N.R., Parkhill, J.: Genetic flux over time in the salmonella lineage. Genome biology 8(6), 100 (2007)

53 Hommais, F., Krin, E., Coppee, J.-Y., Lacroix, C., Yeramian, E., Danchin, A., Bertin, P.: Gade (yhie): a novel activator involved in the response to acid environment in escherichia coli. Microbiology 150(1), 61–72 (2004)

54 Tramonti, A., De Canio, M., De Biase, D.: Gadx/gadw-dependent regulation of the escherichia coli acid fitness island: transcriptional control at the gady–gadw divergent promoters and identification of four novel 42 bp gadx/gadw-specific binding sites. Molecular microbiology 70(4), 965–982 (2008)

55 Wyckoff, E.E., Duncan, D., Torres, A.G., Mills, M., Maase, K., Payne, S.M.: Structure of the shigella dysenteriae haem transport locus and its phylogenetic distribution in enteric bacteria. Molecular microbiology 28(6), 1139–1152 (1998)

56 Bergholz, T.M., Tarr, C.L., Christensen, L.M., Betting, D.J., Whittam, T.S.: Recent gene conversions between duplicated glutamate decarboxylase genes (gada and gadb) in pathogenic escherichia coli. Molecular biology and evolution 24(10), 2323–2333 (2007)

57 Grant, M.A., Weagant, S.D., Feng, P.: Glutamate decarboxylase genes as a prescreening marker for detection of pathogenic escherichia coligroups. Applied and environmental microbiology 67(7), 3110–3114 (2001)

58 Tillman, G., Simmons, M., Wasilenko, J., Narang, N., Cray, W., Bodeis-Jones, S., Martin, G., Gaines, S., Seal, B.: Development of a real-time pcr for escherichia coli based on gade, an acid response regulatory gene. Letters in applied microbiology 60(2), 196–202 (2015)

